# C2c2 is a single-component programmable RNA-guided RNA-targeting CRISPR effector

**DOI:** 10.1101/054742

**Authors:** Omar O. Abudayyeh, Jonathan S. Gootenberg, Silvana Konermann, Julia Joung, Ian M. Slaymaker, David B.T. Cox, Sergey Shmakov, Kira S. Makarova, Ekaterina Semenova, Leonid Minakhin, Konstantin Severinov, Aviv Regev, Eric S. Lander, Eugene V. Koonin, Feng Zhang

## Abstract

The CRISPR-Cas adaptive immune system defends microbes against foreign genetic elements via DNA or RNA-DNA interference. We characterize the Class 2 type VI-A CRISPR-Cas effector C2c2 and demonstrate its RNA-guided RNase function. C2c2 from the bacterium *Leptotrichia shahii* provides interference against RNA phage.*In vitro* biochemical analysis show that C2c2 is guided by a single crRNA and can be programmed to cleave ssRNA targets carrying complementary protospacers. In bacteria, C2c2 can be programmed to knock down specific mRNAs. Cleavage is mediated by catalytic residues in the two conserved HEPN domains, mutations in which generate catalytically inactive RNA-binding proteins. These results broaden our understanding of CRISPR-Cas systems and suggest that C2c2 can be used to develop new RNA-targeting tools.

---

Almost all archaea and about half of bacteria possess Clustered Regularly Interspaced Short Palindromic Repeats and CRISPR-associated genes (CRISPR-Cas) adaptive immune systems (Makarova et al., 2011; Makarova et al., 2015), which protect microbes from viruses and other invading DNA through three steps: (i) adaptation, i.e., insertion of foreign nucleic acid segments (spacers) into the CRISPR array in between pairs of direct repeats (DRs), (ii) transcription and processing of the CRISPR array to produce mature CRISPR RNAs (crRNAs), and (iii) interference, whereby Cas enzymes are guided by the crRNAs to target and cleave cognate sequences in the respective invader genomes (Heler et al., 2015b; van der Oost et al., 2009; Wright et al., 2016. All CRISPR-Cas systems characterized to date follow these three steps, although the mechanistic implementation and proteins involved in these processes display extensive diversity.

The CRISPR-Cas systems are broadly divided into two classes on the basis of the architecture of the interference module: Class 1 systems rely on multi-subunit protein complexes whereas Class 2 systems utilize single effector proteins (Makarova et al., 2015). Within these two classes, types and subtypes are delineated according to the presence of distinct signature genes, protein sequence conservation, and organization of the respective genomic loci. Class 1 systems include type I,where interference is achieved through assembly of multiple Cas proteins into the Cascade complex, and type III systems, which rely on either the Csm (type III-A/D) or Cmr (Type III-B/C) effector complexes which are distantly related to the Cascade (Brouns et al., 2008; Hale et al., 2009; Jackson et al., 2014; Makarova et al., 2015; Marraffini and Sontheimer, 2008; Mulepati et al., 2014; Sinkunas et al., 2013).

Class 2 CRISPR systems comprise type II, characterized by the single-component effector protein Cas9 (Barrangou et al., 2007; Deltcheva et al., 2011; Garneau et al., 2010; Gasiunas et al., 2012; Jinek et al., 2012; Sapranauskas et al., 2011), which contains RuvC and HNH nuclease domains, and type V systems, which utilize single RuvC domain-containing effectors such as Cpf1 (Zetsche et al., 2015), C2c1, and C2c3 (Shmakov et al., 2015). All functionally characterized systems, to date, have been reported to target DNA, and only the multi-component type III-A and III-B systems additionally target RNA Hale et al., 2012; Hale et al., 2009; Jiang et al., 2016; Samai et al., 2015; Staals et al., 2013; Staals et al., 2014; Tamulaitis et al., 2014. However, the putative Class 2 type VI system is characterized by the presence of the single effector protein C2c2, which lacks homology to any known DNA nuclease domain but contains two Higher Eukaryotes and Prokaryotes Nucleotide-binding (HEPN) domains (Shmakov et al., 2015). Given that all functionally characterized HEPN domains are RNases (Anantharaman et al., 2013), there is a possibility that C2c2 functions solely as an RNA-guided RNA-targeting CRISPR effector.

HEPN domains are also found in other Cas proteins. Csm6, a component of type III-A systems, and the homologous protein Csx1, in type III-B systems, each contain a single HEPN domain and have been biochemically characterized as ssRNA-specific endoribonucleases (Jiang et al., 2016; Niewoehner and Jinek, 2016; Sheppard et al., 2016). In addition, type III systems contain complexes of other Cas enzymes that bind and cleave ssRNA through acidic residues associated with RNA-recognition motif (RRM) domains. These complexes (Cas10-Csm in type III-A and Cmr in type III-B) carry out RNA-guided co-transcriptional cleavage of mRNA in concert with DNA target cleavage (Deng et al., 2013; Goldberg et al., 2014; Samai et al., 2015). In contrast, the roles of Csm6 and Csx1, which cleave their targets with little specificity, are less clear, although in some cases, RNA cleavage by Csm6 apparently serves as a second line of defense when DNA targeting fails (Jiang et al., 2016). Additionally Csm6 and Csx1 have to dimerize to form a composite active site (Kim et al., 2013; Niewoehner and Jinek, 2016; Sheppard et al., 2016), but C2c2 contains two HEPN domains, suggesting that it functions as a monomeric endoribonuclease.

As is common with Class 2 systems, type VI systems are simply organized. In particular, the type VI locus in *Leptotrichia shahii* contains Cas1, Cas2, C2c2 and a CRISPR array, which is expressed and processed into mature crRNAs (Shmakov et al., 2015). In all CRISPR-Cas systems characterized to date, Cas1 and Cas2 are exclusively involved in spacer acquisition (Datsenko et al., 2012; Diez-Villasenor et al., 2013; Heler et al., 2015a; Nunez et al., 2014; Nunez et al., 2015; Yosef et al., 2012), suggesting that C2c2 is the sole effector protein which utilizes a crRNA guide to achieve interference, likely targeting RNA.

## Reconstitution of the L.*shahii* C2c2 locus in *Escherichia coli* confers RNA-guided immunity

We explored whether LshC2c2 could confer immunity to MS2 (Tamulaitis et al., 2014), a lytic single-stranded (ss) RNA phage, without DNA intermediates in its life cycle, that infects *E. coli*. We constructed a low-copy plasmid carrying the entire LshC2c2 locus (pLshC2c2) to allow for heterologous reconstitution in *E. coli* (Figure S1A). Because expressed mature crRNAs from the LshC2c2 locus have a maximum spacer length of 28nt (Figure S1A) (Shmakov et al., 2015), we tiled all possible 28-nt target sites in the MS2 phage genome (Figure 1A). This resulted in a library of 3,473 spacer sequences (along with 588 non-targeting guides designed to have a Levenshtein distance of ≥8 with respect to the MS2 and *E. coli* genomes) which we inserted between pLshC2c2 direct repeats (DRs). After transformation in of this construct into *E. coli*, we infected cells with varying dilutions of MS2(10^−1^, 10^−3^, and 10^−5^) and analyzed surviving cells to determine the spacer sequences carried by cells that survived the infection. Cells carrying spacers that confer robust interference against MS2 are expected to proliferate faster than those that lack such sequences. Following growth for 16 hours, we identified a number of spacers that were consistently enriched across three independent infection replicas in both the 10^−1^ and 10^−3^ dilution conditions, suggesting that they enabled interference against MS2. Specifically, 152 and 144 spacers showed >0.8 log_2_fold enrichment in all three replicates for the 10^−1^ and 10^−3^ phage dilutions, respectively; of these two groups of top enriched spacers, 75 are shared (Figs. 1B, S2A-G, table S1). Additionally, no non-targeting guides were found to be consistently enriched among the three 10^−1^, 10^−3^, or 10^−5^ phage replicates (Figure S2D, G). We also analyzed the flanking regions of protospacers on the MS2 genome corresponding to the enriched spacers and found that spacers with a G immediately flanking the 3’ end of the protospacer were less fit relative to all other nucleotides at this position (i.e. A,U,or C), suggesting that the 3′ protospacer flanking site (PFS) affects the efficacy of C2c2-mediated targeting (Figs. 1C, S2E-F, S3). Although the PFS is adjacent to the protospacer target, we chose not to use the commonly used protospacer adjacent motif (PAM) nomenclature as it has come to connote a sequence used in self vs. non-self differentiation (Marraffini and Sontheimer,2010), which is irrelevant in a RNA-targeting system. It is worth noting that the avoidance of G by C2c2 echo the absence of PAMs applicable to other RNA-targeting CRISPR systems and effector proteins (Hale et al., 2014; Hale et al., 2012; Samai et al., 2015; Staals et al., 2014; Tamulaitis et al., 2014; Zhang et al., 2012).

**Fig. 1.**
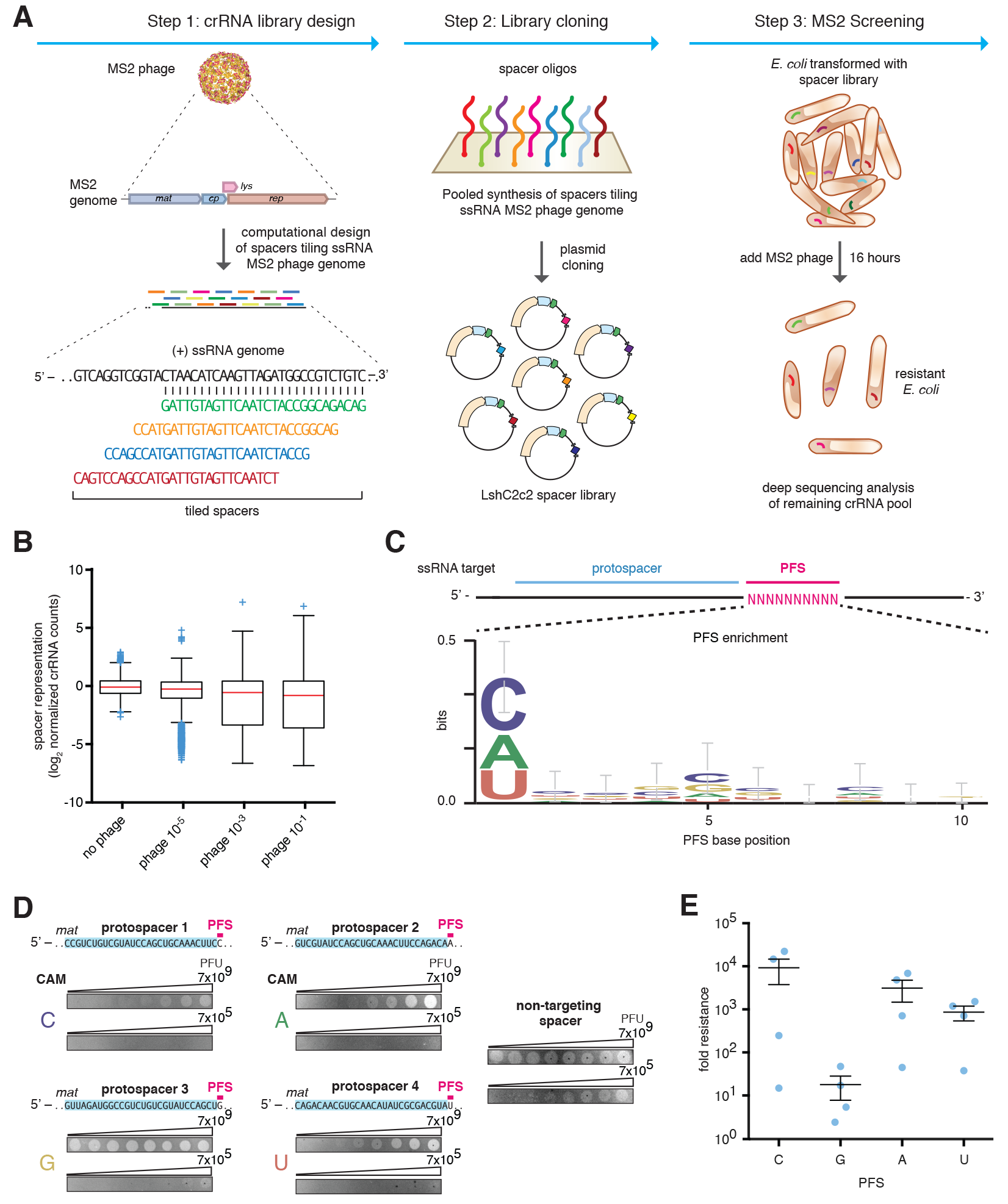
Heterologous expression of the *Leptotrichia shahii* C2c2 locus mediates robust interference of RNA phage in *Escherichia coli*. A) Schematic for the MS2 bacteriophage interference screen. A library consisting of spacers targeting all possible sequences in the MS2 RNA genome was cloned into the LshC2c2 CRISPR array. Cells transformed with the MS2-targeting spacer library were then treated with phage and plated, and surviving cells were harvested. The frequency of spacers was compared to an untreated control (no phage), and enriched spacers from the phage-treated condition were used for the generation of PFS preference logos. B) Box plot showing the distribution of normalized crRNA frequencies for the phage-treated conditions and control screen (no phage) biological replicates (n=3). The box extends from the first to third quartile with whiskers denoting 1.5 times the interquartile range. The mean is indicated by the red horizontal bar. The 10^−1^ and 10^−3^ phage dilution distributions are significantly different than each of the control replicates (****, p<0.0001 by ANOVA with multiple hypothesis correction). C) Sequence logo generated from sequences flanking the 3’ end of protospacers corresponding to enriched spacers in the 10^−3^ phage dilution condition, revealing the presence of a 3’ H PFS (not G). D) Plaque assay used to validate the functional significance of the H PFS in MS2 interference. All protospacers flanked by non-G PFSs exhibited robust phage interference. Spacer were designed to target the MS2 *mat* gene and their sequences are shown above the plaque images; the spacer used in the non-targeting control is not complementary to any sequence in either the *E. coli* or MS2 genome. Phage spots were applied as series of half-log dilutions. E) Quantitation of MS2 plaque assay validating the H (non-G) PFS preference. 4 MS2-targeting spacers were designed for each PFS. Each point on the scatter plot represents the average of three biological replicates and corresponds to a single spacer. Bars indicate the mean of 4 spacers for each PFS and standard error (s.e.m).

The fact that only ~5% of crRNAs are enriched may reflect other factors influencing interference activity, such as accessibility of the target site that might be affected by RNA binding proteins or secondary structure. In agreement with this hypothesis, the enriched spacers tend to cluster into regions of strong interference where they are closer to each other than one would expect by random chance (Figure S3F-G).

To validate the interference activity of the enriched spacers, we individually cloned four top-enriched spacers into pLshC2c2 CRISPR arrays and observed a 3- to 4-log_10_ reduction in plaque formation, consistent with the level of enrichment observed in the screen (Figs 1B, S4). We cloned sixteen guides targeting distinct regions of the MS2 *mat* gene (4 guides per possible singlenucleotide PFS). All 16 crRNAs mediated MS2 interference, although higher levels of resistance were observed for the C, A, and U PFS-targeting guides (Figs. 1D, 1E, S5), indicating that C2c2 can be effectively retargeted in a crRNA-dependent fashion to sites within the MS2 genome.

To further validate the observed PFS preference with an alternate approach, we designed a protospacer site in the pUC19 plasmid at the 5’ end of the ß-lactamase mRNA, which encodes ampicillin resistance in *E. coli*, flanked by five randomized nucleotides at the 3’ end. Significant depletion and enrichment was observed for the LshC2c2 locus (****, p<0.0001) compared to the pACYC184 controls (Figure S6A). Analysis of the depleted PFS sequences confirmed the presence of a PFS preference of H (Figure S6B).

## C2c2 is a single-effector endoRNase mediating ssRNA cleavage with a single crRNA guide

We purified the LshC2c2 protein (Figure S7) and assayed its ability to cleave an *in vitro* transcribed 173-nt ssRNA target (Figs. 2A, S8) containing a C PFS (ssRNA target 1 with protospacer 14). Mature LshC2c2 crRNAs contain a 28-nt direct repeat (DR) and a 28 nt spacer (Figure S1A)(Shmakov et al., 2015). We therefore generated an *in-vitro*-transcribed crRNA with a 28-nt spacer complementary to protospacer 14 on ssRNA target 1. LshC2c2 efficiently cleaved ssRNA in a Mg^2+^- and crRNA-dependent manner (Figs. 2B, S9). We then annealed complementary RNA oligos to regions flanking the crRNA target site. This partially double-stranded RNA substrate was not cleaved by LshC2c2, suggesting it is specific for ssRNA (figs. S10A-B).

**Figure. 2.**
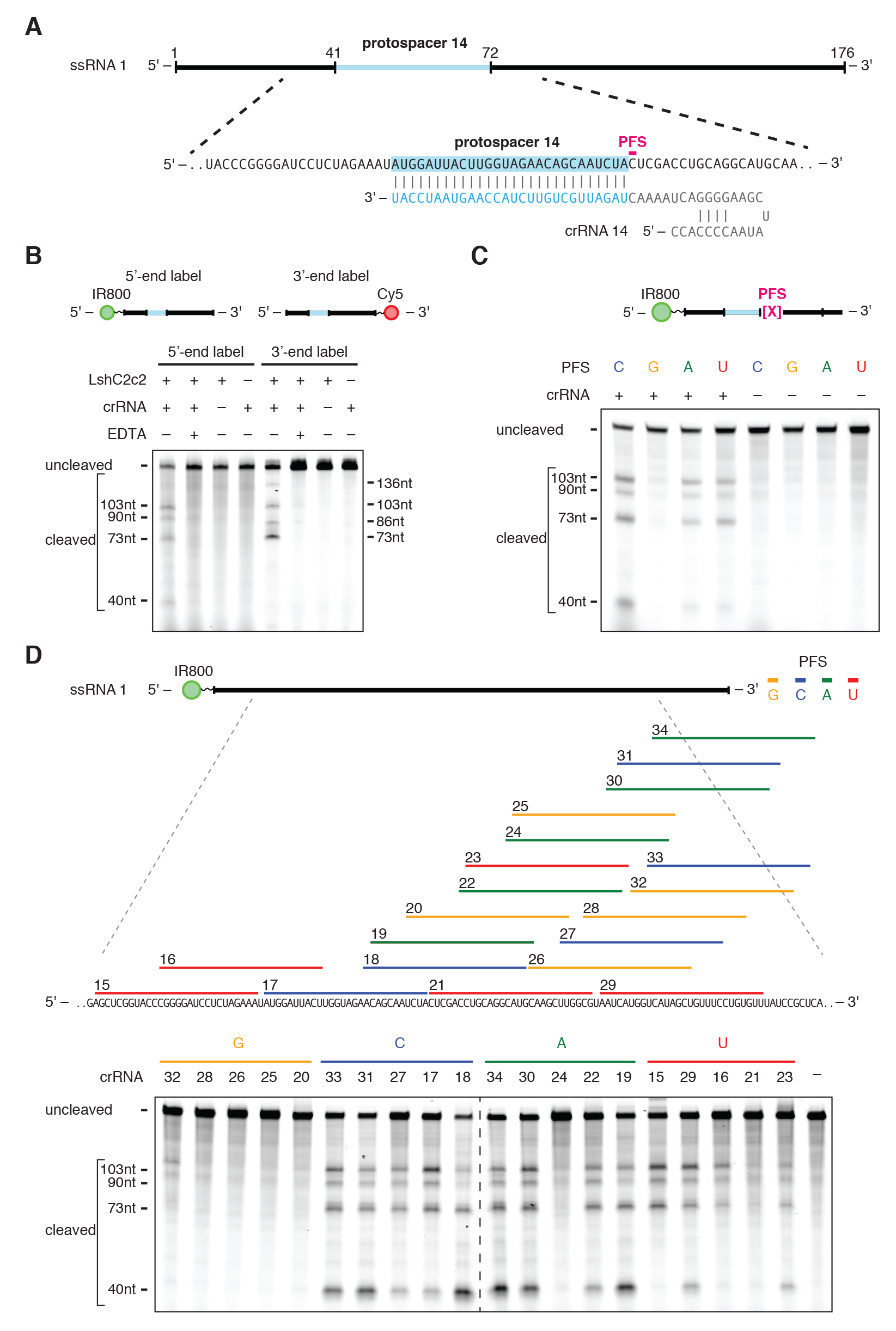
LshC2c2 and crRNA mediate RNA-guided ssRNA cleavage. A) Schematic of the ssRNA substrate being targeted by the crRNA. The protospacer region is highlighted in blue and the PFS is indicated by the magenta bar. B) A denaturing gel demonstrating crRNA-mediated ssRNA cleavage by LshC2c2 after 1 hour of incubation. The ssRNA target is either 5’ labeled with IRDye 800 or 3’ labeled with Cy5. Cleavage requires the presence of the crRNA and is abolished by addition of EDTA. Four cleavage sites are observed. Reported band lengths are matched from RNA sequencing. C) A denaturing gel demonstrating the requirement for an H PFS (not G) after 3 hours of incubation. Four ssRNA substrates that are identical except for the PFS (indicated by the magenta X in the schematic) were used for the *in vitro* cleavage reactions. ssRNA cleavage activity is dependent on the nucleotide immediately 3’ of the target site. Reported band lengths are matched from RNA sequencing. D) Schematic showing five protospacers for each PFS on the ssRNA target (top). Denaturing gel showing crRNA-guided ssRNA cleavage activity after 1 hour of incubation. crRNAs correspond to protospacer numbering. Reported band lengths are matched from RNA sequencing.

We tested the sequence constraints of RNA cleavage by LshC2c2 with additional crRNAs complementary to ssRNA target 1 where protospacer 14 is preceded by each PFS variant.The results of this experiment confirmed the preference for C, A, and U PFSs, with little cleavage activity detected for the G PFS target (Figure 2C). Additionally, we designed 5 crRNAs for each possible PFS (20 total) across the ssRNA target 1 and evaluated cleavage activity for LshC2c2 paired with each of these crRNAs. As expected, we observed less cleavage activity for G PFS-targetingcrRNAs compared to other crRNAs tested (Figure 2D).

We then generated a dsDNA plasmid library with protospacer 14 flanked by 7 random nucleotides to account for any PFS preference. When incubated with LshC2c2 protein and a crRNA complementary to protospacer 14, no cleavage of the dsDNA plasmid library was observed (Figure S10C). We also did not observe cleavage when targeting a ssDNA version of ssRNA target 1 (Figure S10D). To rule out co-transcriptional DNA cleavage, which has been observed in type III CRISPR-Cas systems(Samai et al., 2015), we recapitulated the *E. coli* RNA polymerase co-transcriptional cleavage assay (Samai et al., 2015)(Figure S11A) expressing ssRNA target 1 from a DNA substrate. This assay of purified LshC2c2 and crRNA targeting ssRNA target 1 did not show any DNA cleavage (Figure S11B). Together, these results indicate that C2c2 cleaves specific ssRNA sites directed by the target complementarity encoded in the crRNA, with a H PFS preference.

## C2c2 cleavage depends on local target sequence and secondary structure

Given that C2c2 did not efficiently cleave dsRNA substrates and that ssRNA can form complex secondary structures, we reasoned that cleavage by C2c2 might be affected by secondary structure of the ssRNA target. Indeed, after tiling ssRNA target 1 with different crRNAs (Figure 2D), we observed the same cleavage pattern regardless of the crRNA position along the target RNA. This observation suggests that the crRNA-dependent cleavage pattern was determined by features of the target sequence rather than the distance from the binding site. We hypothesized that the LshC2c2-crRNA complex binds the target and cleaves exposed regions of ssRNA within the secondary structure elements, with potential preference for certain nucleotides.

In agreement with this hypothesis, cleavage of three ssRNA targets with different sequences flanking identical 28-nt protospacers resulted in three distinct patterns of cleavage (Figure 3A). RNA-sequencing of the cleavage products for the three targets revealed that cleavage sites mainly localized to uracil-rich regions of ssRNA or ssRNA-dsRNA junctions within the *in silico* predicted co-folds of the target sequence with the crRNA (Figs. 3B-C, S12A-D). To test whether the LshC2c2-crRNA complex prefers cleavage at uracils, we analyzed the cleavage efficiencies of homopolymeric RNA targets (a 28-nt protospacer extended with 120 As or Us regularly interspaced by single bases of G or C to enable oligo synthesis) and found that LshC2c2 preferentially cleaved the uracil target compared to adenine (figs. S12E, S12F). We then tested cleavage of a modified version of ssRNA 4 which had its main site of cleavage, a loop, replaced with each of the four possible homopolymers and found that cleavage only occurred at the uracil homopolymer loop (Figure S12G). To further test whether cleavage was occurring at uracil residues, we mutated single uracil residues in ssRNA 1 that showed cleavage in the RNA-sequencing (Figure 3B) to adenines. This experiment showed that, by mutating each uracil residue, we could modulate the presence of a single cleavage band, consistent with LshC2c2 cleaving at uracil residues in ssRNA regions (Figure 3D).

**Fig. 3.**
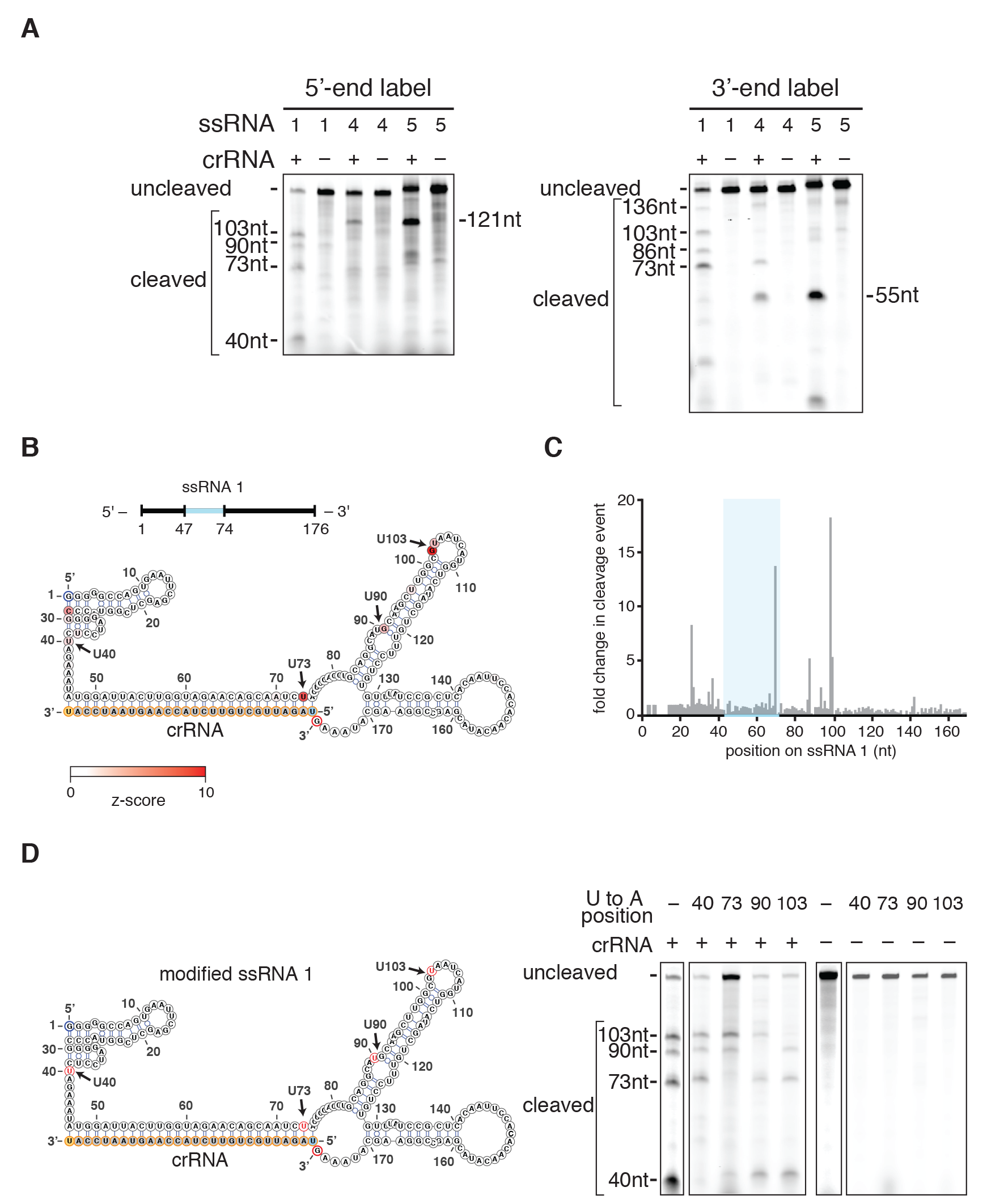
C2c2 cleavage sites are determined by secondary structure and sequence of the target RNA. A) Denaturing gel showing C2c2-crRNA-mediated cleavage after 3 hours of incubation of three non-homopolymeric ssRNA targets (1,4,5; black, blue and green on figs 3B-C and S12A-D respectively) that share the same protospacer but are flanked by different sequences. Despite identical protospacers, different flanking sequences resulted in different cleavage patterns. Reported band lengths are matched from RNA sequencing. B) The cleavage sites of non-homopolymer ssRNA target 1 were mapped with RNA-sequencing of the cleavage products. The frequency of cleavage at each base is colored according to the z-score and shown on the predicted crRNA-ssRNA co-fold secondary structure. Fragments used to generate the frequency analysis contained the complete 5’ end. The 5’ and 3’ end of the ssRNA target are indicated by blue and red outlines, on the ssRNA and secondary structure, respectively. The 5’ and 3’ end of the spacer (outlined in yellow) is indicated by the blue and orange residues highlighted respectively. The crRNA nucleotides are highlighted in orange. C) Plot of the frequencies of cleavage sites for each position of ssRNA target 1 for all reads that begin at the 5’ end. The protospacer is indicated by the blue highlighted region. D) Schematic of a modified ssRNA 1 target showing sites (*red*) of single U to A flips (*left*). Denaturing gel showing C2c2-crRNA mediated cleavage of each of these single nucleotide variants after 3 hours of incubation (*right*). Reported band lengths are matched from RNA sequencing.

## The HEPN domains of C2c2 mediate RNA-guided ssRNA-cleavage

Bioinformatic analysis of C2c2 has suggested that the HEPN domains are likely to be responsible for the observed catalytic activity (Shmakov et al., 2015). Each of the two HEPN domains of C2c2 contains a dyad of conserved arginine and histidine residues (Figure 4A), in agreement with the catalytic mechanism of the HEPN endoRNAse (Anantharaman et al., 2013); Niewoehner and Jinek, 2016; Sheppard et al., 2016). We mutated each of these putative catalytic residues separately to alanine (R597A, H602A, R1278A, H1283A) in the LshC2c2 locus plasmids and assayed for MS2 interference. None of the four mutant plasmids were able to protect *E. coli* from phage infection (Figs. 4B, S13).

**Fig. 4.**
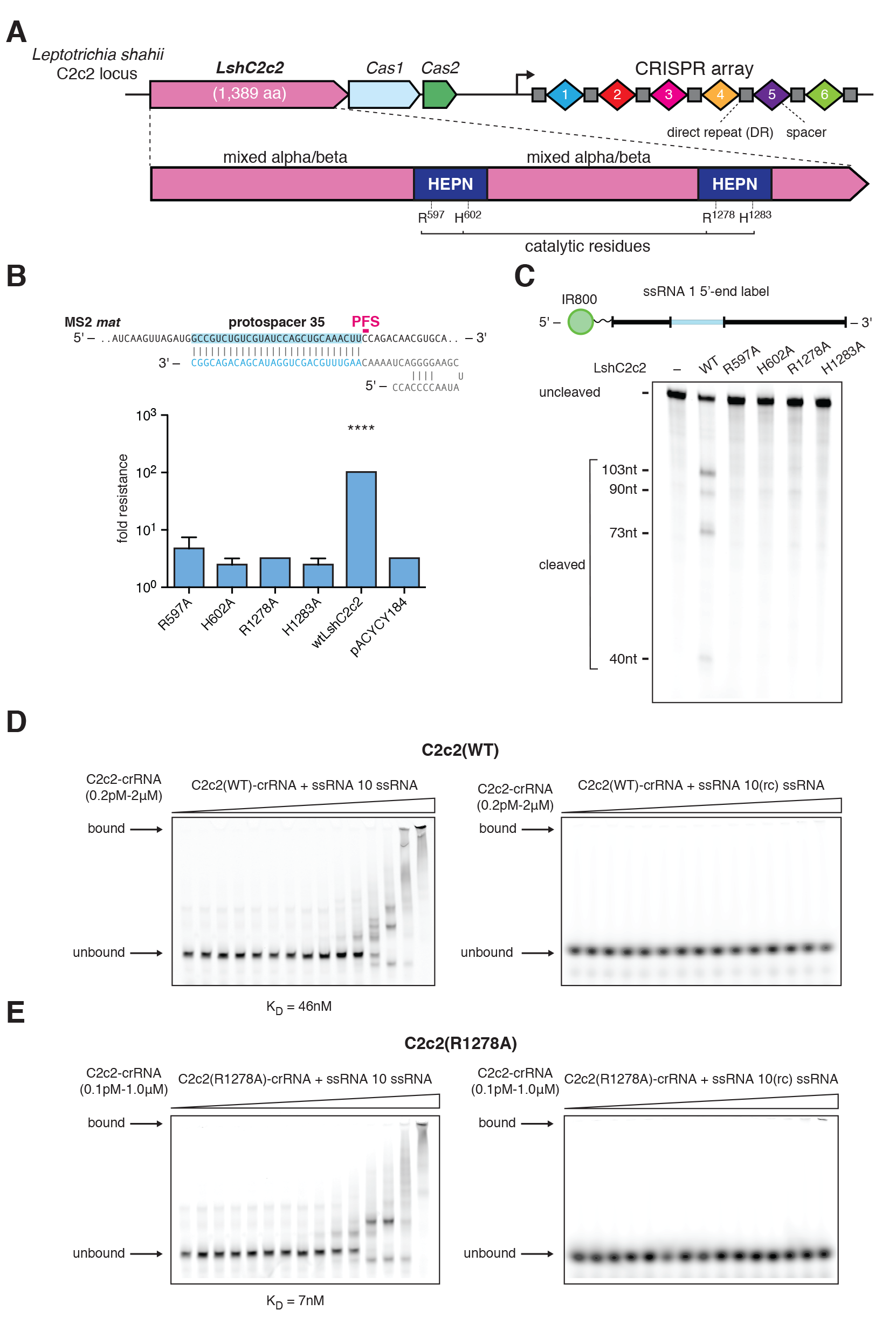
The two HEPN domains of C2c2 are necessary for crRNA-guided ssRNA cleavage but not for binding. A) Schematic of the *LshC2c2* locus and the domain organization of the LshC2c2 protein, showing conserved residues in HEPN domains (dark blue). B) Quantification of MS2 plaque assay with HEPN catalytic residue mutants. For each mutant, the same crRNA targeting protospacer 35 was used. (n=3 biological replicates, ****, p<0.0001 compared to pACYC184 by t-test. Bars represent mean ± s.e.m.) C) Denaturing gel showing conserved residues of the HEPN motif, indicated as catalytic residues in panel A, are necessary for crRNA-guided ssRNA target 1 cleavage after 3 hours of incubation. Reported band lengths are matched from RNA sequencing. D) Electrophoretic mobility shift assay (EMSA) evaluating affinity of the wild type LshC2c2-crRNA complex against a targeted (left) and a non-targeted (right) ssRNA substrate. The non-targeted ssRNA substrate is the reverse-complement of the targeted ssRNA 10.EDTA is supplemented to reaction condition to reduce any cleavage activity. E) Electrophoretic mobility shift assay with LshC2c2(R1278A)-crRNA complex against on-target ssRNA 10 and non-targeting ssRNA (same substrate sequences as in D)

We purified the four single-point mutant proteins and assayed their ability to cleave 5’-end-labeled ssRNA target 1 (Figure 4C). In agreement with our *in vivo* results, all four mutations abolished cleavage activity. The inability of either of the two wild-type HEPN domains to compensate for inactivation of the other implies cooperation between the two domains. These results agree with observations that several bacterial and eukaryotic single-HEPN proteins function as dimers (Kozlov et al., 2011; Niewoehner and Jinek, 2016; Sheppard et al., 2016).

Catalytically inactive variants of Cas9 retain target DNA binding, allowing for the creation of programmable DNA-binding proteins (Gasiunas et al., 2012; Jinek et al., 2012). Electrophoretic mobility shift assays (EMSA) on both the wild-type (Figure 4D) and R1278A mutant LshC2c2 (Figure 4E) in complex with crRNA showed the wild-type LshC2c2 complex binding strongly (K_D_ ~ 46 nM, Figure S14A) and specifically to 5’-end-labeled ssRNA target 10 but not to the 5’-end-labeled non-target ssRNA (the reverse complement of ssRNA target 10). The R1278A mutant C2c2 complex showed even stronger (K_D_ ~ 7 nM, Figure S14B) specific binding, indicating that this HEPN mutation results in a catalytically inactive, RNA-programmable RNA-binding protein. The LshC2c2 protein or crRNA alone showed reduced levels of target affinity, as expected (Figure S14C-E). Additionally, no specific binding of LshC2c2-crRNA complex to ssDNA was observed (Figure S15).

These results demonstrate that C2c2 cleaves RNA via a catalytic mechanism distinct from other known CRISPR-associated RNases. In particular, the type III Csm and Cmr multiprotein complexes rely on acidic residues of RRM domains for catalysis, whereas C2c2 achieves RNA cleavage through the conserved basic residues of its two HEPN domains.

## Sequence and structural requirements of C2c2 crRNA

Similar to the type V-B (Cpf1) systems (Zetsche et al., 2015), the LshC2c2 crRNA contains a single stem loop in the direct repeat (DR), suggesting that the secondary structure of the crRNA could facilitate interaction with LshC2c2. We thus investigated the length requirements of the spacer sequence for ssRNA cleavage and found that LshC2c2 requires spacers of at least 22 nt length to efficiently cleave ssRNA target 1 (Figure S16A). The stem-loop structure of the crRNA is also critical for ssRNA cleavage, because DR truncations that disturbed the stem loop abrogated target cleavage (Figure S16B). Thus, a DR longer than 24 nt is required to maintain the stem loop necessary for LshC2c2 to mediate ssRNA cleavage.

Single base pair inversions in the stem that preserved the stem structure did not affect the activity of the LshC2c2 complex. In contrast, inverting all four G-C pairs in the stem eliminated the cleavage despite maintaining the duplex structure (Figure S17A). Other perturbations, such as those that introduced kinks and reduced or increased base-pairing in the stem, also eliminated or drastically suppressed cleavage. This suggests that the crRNA stem length is important for complex formation and activity (Figure S17A). We also found that loop deletions eliminated cleavage, whereas insertions and substitutions mostly maintained some level of cleavage activity (Figure S17B). In contrast, nearly all substitutions or deletions in the region 3’ to the DR prevented cleavage by LshC2c2 (fig S18). Together, these results demonstrate that LshC2c2 recognizes structural characteristics of its cognate crRNA but is amenable to loop insertions and most tested base substitutions outside of the 3’ DR region. These results have implications for the future application of C2c2-based tools that require guide engineering for recruitment of effectors or modulation of activity (Dahlman et al., 2015; Kiani et al., 2015; Konermann et al., 2015).

## C2c2 cleavage is sensitive to double mismatches in the crRNA-target duplex

We tested the sensitivity of the LshC2c2 system to single mismatches between the crRNA guide and target RNA by mutating single bases across the spacer to the respective complementary bases (e.g., A to U). We then quantified plaque formation with these mismatched spacers in the MS2 infection assay and found that C2c2 was fully tolerant to single mismatches across the spacer as such mismatched spacers interfered with phage propagation with similar efficiency as fully matched spacers (figs. S19A, S20). However, when we introduced consecutive double substitutions in the spacer, we found a ~3 log_10_-fold reduction in the protection for mismatches in the center, but not at the 5’-or 3’-end, of the crRNA (figs. 19B, S20). This observation suggests the presence of a mismatch-sensitive “seed region” in the center of the crRNA-target duplex.

We generated a set of *in vitro* transcribed crRNAs with mismatches similarly positioned across the spacer region. When incubated with LshC2c2 protein, all single mismatched crRNA supported cleavage (Figure S19C), in agreement with our *in vivo* findings. When tested with a set of consecutive and non-consecutive double mutant crRNAs, LshC2c2 was unable to cleave the target RNA if the mismatches were positioned in the center, but not at the 5’-or 3’-end of the crRNA (Figure S19D, S21A), further supporting the existence of a central seed region. Additionally, no cleavage activity was observed with crRNAs containing consecutive triple mismatches in the seed region (Figure S21B).

## C2c2 can be reprogrammed to mediate specific mRNA knockdown *in vivo*

Given the ability of C2c2 to cleave target ssRNA in a crRNA sequence-specific manner, we tested whether LshC2c2 could be reprogrammed to degrade selected non-phage ssRNA targets, and particularly mRNAs, *in vivo*. We co-transformed *E.coli* with a plasmid encoding LshC2c2 and a crRNA targeting the mRNA of red fluorescent protein (RFP) as well as a compatible plasmid expressing RFP (Figure 5A). For OD-matched samples, we observed an approximately 20% to 92% decrease in RFP positive cells for crRNAs targeting protospacers flanked by C, A, or U PFSs (Figure 5B,C). As a control, we tested crRNAs containing reverse complements (targeting the dsDNA plasmid) of the top performing RFP mRNA-targeting spacers. As expected, we observed no decrease in RFP fluorescence by these crRNAs (Figure 5B). We also confirmed that mutation of the catalytic arginine residues in either HEPN domain to alanine precluded RFP knockdown (Figure S22). Thus, C2c2 is capable of general retargeting to arbitrary ssRNA substrates, governed exclusively by predictable nucleic-acid interactions.

**Fig. 5.**
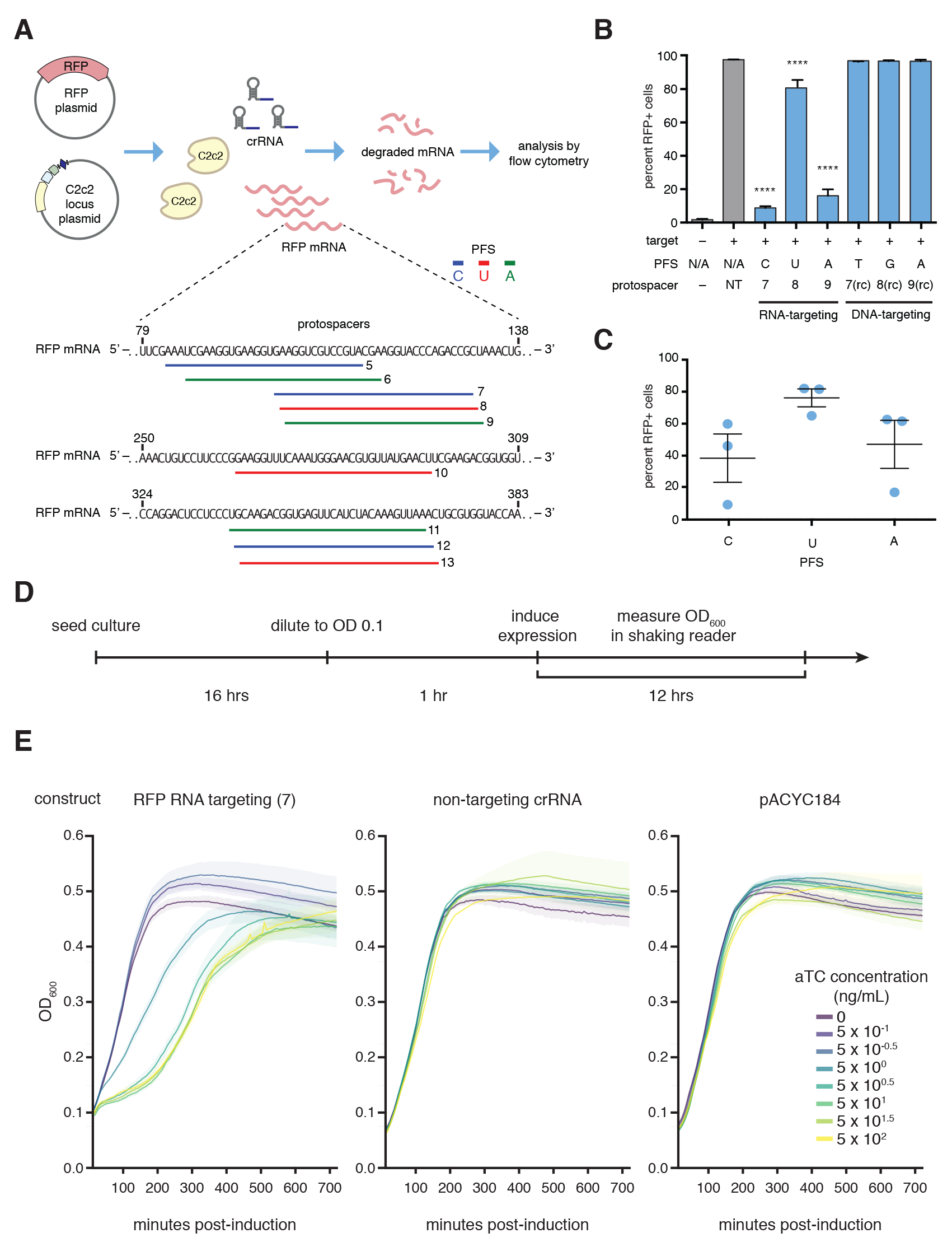
RFP mRNA knockdown by retargeting LshC2c2. A) Schematic showing crRNA-guided knockdown of RFP in *E. coli* heterologously expressing the LshC2c2 locus. Three RFP-targeting spacers were selected for each non-G PFS and each protospacer on the RFP mRNA is numbered. B) RFP mRNA-targeting spacers effected RFP knockdown whereas DNA-targeting spacers (targeting the non-coding strand of the RFP gene on the expression plasmid, indicated as “rc” spacers) did not affect RFP expression. (n=3 biological replicates, ****, p<0.0001 compared to non-targeting guide by ANOVA with multiple hypothesis correction. Bars represent mean ± s.e.m) C) Quantification of RFP knockdown in *E.coli.*Three spacers each targeting C,U,or A PFS-flanking protospacers (9 spacers, numbered 5-13 as indicated in panel (A)) in the RFP mRNA were introduced and RFP expression was measured by flowcytometry. Each point on the scatter plot represents the average of three biological replicates and corresponds to a single spacer. Bars indicate the mean of 3 spacers for each PFS and errors bars are shown as the s.e.m. D) Timeline of *E. coli* growth assay. E) Effect of RFP mRNA targeting on the growth rate of *E. coli* transformed with an inducible RFP expression plasmid as well as the LshC2c2 locus with non-targeting, RNA targeting (spacer complementary to the RFP mRNA or RFP gene coding strand), and pACYC control plasmid at different anhydrotetracycline (aTc) concentrations.

When we examined the growth of cells carrying the RFP-targeting spacer with the greatest level of RFP knockdown, we noted that the growth rate of these bacteria was substantially reduced (Figure 5A, spacer 7). We investigated whether the effect on growth was mediated by the RFP mRNA-targeting activity of LshC2c2 by introducing an inducible-RFP plasmid and an RFP-targeting LshC2c2 locus into *E.coli.* Upon induction of RFP transcription, cells with RFP knockdown showed substantial growth suppression, not observed in non-targeting controls (Figure 5D, E). This growth restriction was dependent on the level of the RFP mRNA, as controlled by the concentration of the inducer anhydrotetracycline. In contrast, in the absence of RFP transcription, we did not observe any growth restriction nor did we observe any transcription-dependent DNA targeting in our biochemical experiment (Figure S11). These results indicate that RNA-targeting is likely the primary driver of this growth restriction phenotype. We therefore surmised that, in addition to the cleavage of the target RNA, C2c2 CRISPR systems might prevent virus reproduction also via non-specific cleavage of cellular mRNAs, causing programmed cell death (PCD) or dormancy (Hayes and Van Melderen, 2011; Makarova et al., 2009).

## C2c2 cleaves collateral RNA in addition to crRNA-targeted ssRNA

Cas9 and Cpf1 cleave DNA within the crRNA-target heteroduplex at defined positions, reverting to an inactive state after cleavage. In contrast, C2c2 cleaves the target RNA outside of the crRNA binding site at varying distances depending on the flanking sequence, presumably within exposed ssRNA loop regions (Figs. 3B, 3C, S12A-D). This observed flexibility with respect to the cleavage distance led us to test whether cleavage of other, non-target ssRNAs also occurs upon C2c2 target binding and activation. Under this model, the C2c2-crRNA complex, once activated by binding to its target RNA, cleaves the target RNA as well as other RNAs non-specifically. We carried out *in vitro* cleavage reactions that included, in addition to LshC2c2 protein, crRNA and its target RNA, one of four unrelated RNA molecules without any complementarity to the crRNA guide (Figure 6A). These experiments showed that, whereas the LshC2c2-crRNA complex did not mediate cleavage of any of the four collateral RNAs in the absence of the target RNA, all four were efficiently degraded in the presence of the target RNA (Figs. 6B, S23A). Furthermore, R597A and R1278A HEPN mutants were unable to cleave collateral RNA (Figure S23B).

**Fig. 6.**
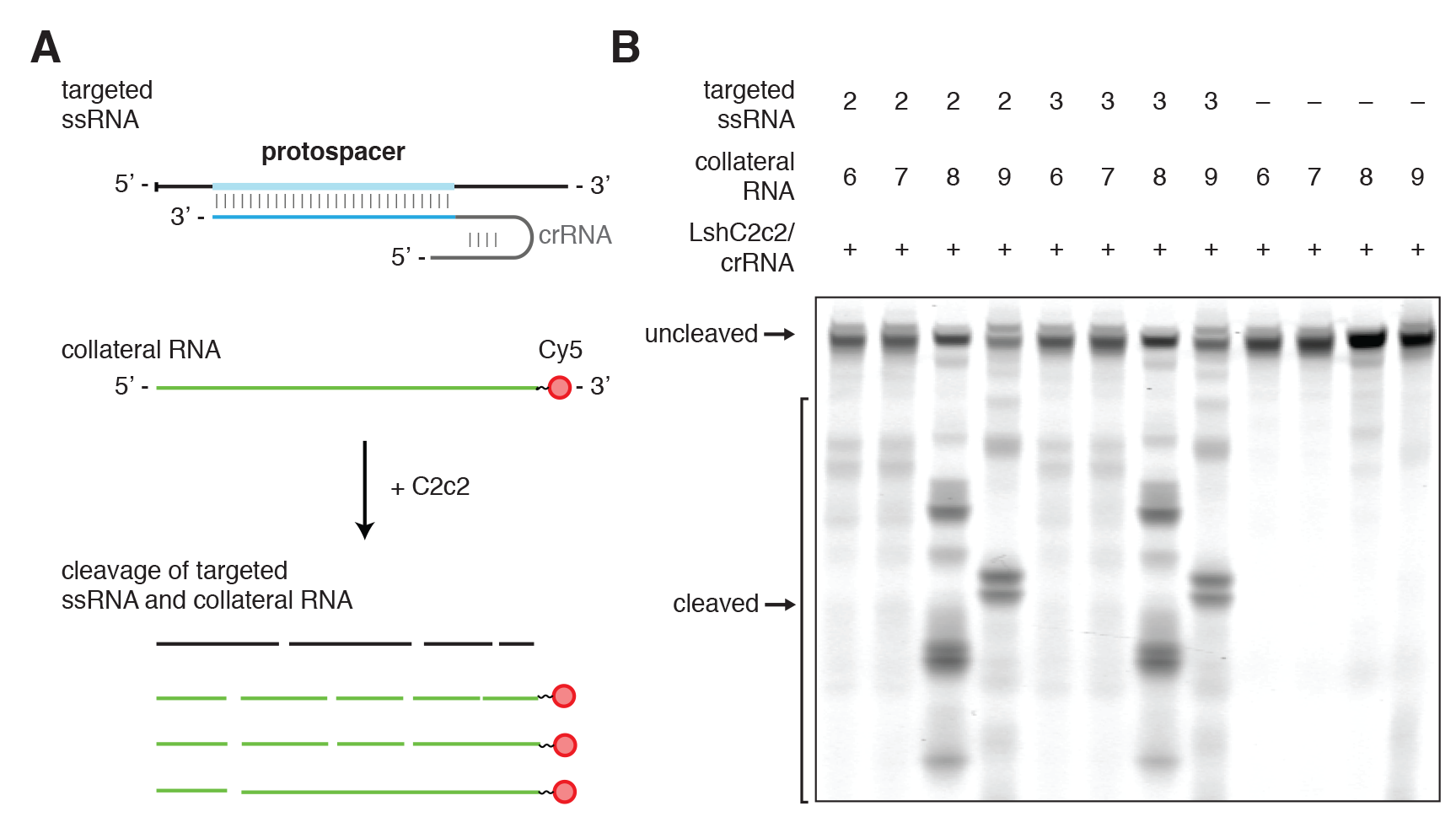
crRNA-guided ssRNA cleavage activates non-specific RNase activity of LshC2c2. A) Schematic of the biochemical assay used to detect crRNA-binding-activated non-specific RNase activity on non-crRNA-targeted collateral RNA molecules. The reaction consists of C2c2 protein, unlabeled crRNA, unlabeled target ssRNA, and a second ssRNA with 3’ fluorescent labeling and is incubated for 3 hours. C2c2-crRNA mediates cleavage of the unlabeled target ssRNA as well as the 3’-end-labeled collateral RNA which has no complementarity to the crRNA. B) Denaturing gel showing non-specific RNase activity against non-targeted ssRNA substrates in the presence of target RNA after 3 hours of incubation. The non-targeted ssRNA substrate is not cleaved in the absence of the crRNA-targeted ssRNA substrate.

To further investigate the collateral cleavage and growth restriction *in vivo*, we hypothesized that if a PFS preference screen for LshC2c2 was performed in a transcribed region on the transformed plasmid, then we should be able to detect the PFS preference due to growth restriction induced by RNA targeting. We designed a protospacer site flanked by five randomized nucleotides at the 3’ end in either a non-transcribed region or in a region transcribed from the *lac* promoter (Figure S24A). The analysis of the depleted and enriched PFS sequences identified a H PFS only in the assay with the transcribed sequence but no discernable motif in the non-transcribed sequence (Figure S24B-C).

These results suggest a HEPN-dependent mechanism whereby C2c2 in a complex with crRNA is activated upon binding to target RNA and subsequently cleaves non-specifically other available ssRNA targets. Such promiscuous RNA cleavage could cause cellular toxicity, resulting in the observed growth rate inhibition. These findings imply that, in addition to their likely role in direct suppression of RNA viruses, type VI CRISPR-Cas systems could function as mediators of a distinct variety of PCD or dormancy induction that is specifically triggered by cognate invader genomes (Figure 7). Under this scenario, dormancy would slow the infection and supply additional time for adaptive immunity. Such a mechanism agrees with the previously proposed coupling of adaptive immunity and PCD during the CRISPR-Cas defensive response (Makarova et al., 2012).

**Fig. 7.**
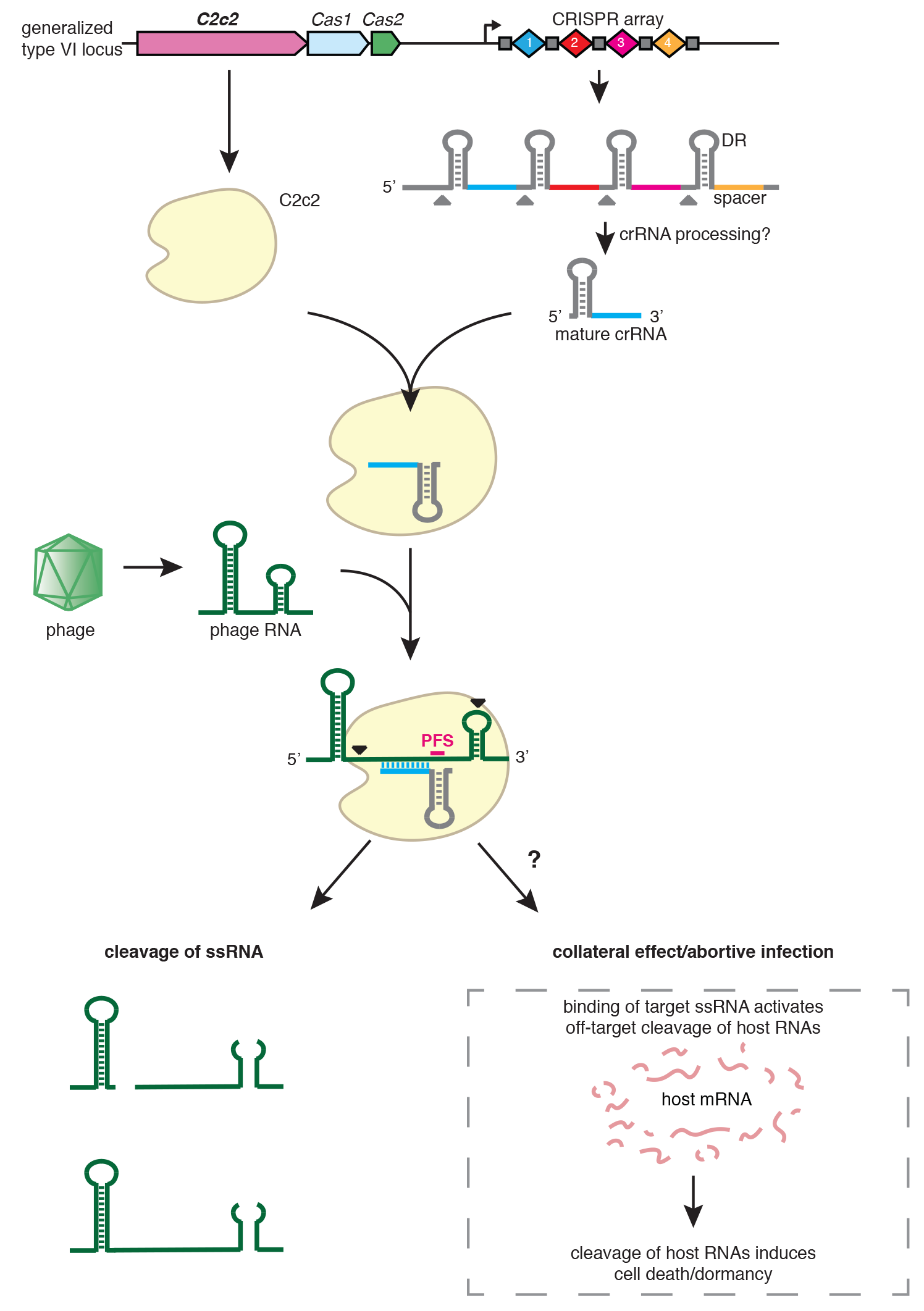
C2c2 as a putative RNA-targeting prokaryotic immune system. The C2c2-crRNA complex recognizes target RNA via base pairing with the cognate protospacer and cleaves the target RNA. In addition, binding of the target RNA byC2c2-crRNA activates a non-specific RNase activity which may lead to promiscuous cleavage of RNAs without complementarity to the crRNA guide sequence. Through this nonspecific RNase activity, C2c2 may also cause abortive infection via programmed cell death or dormancy induction.

## Conclusions

In summary, the Class 2 type VI effector protein C2c2 is an RNA-guided RNase that can be efficiently programmed to degrade any ssRNA by specifying a 28-nt sequence on the crRNA (Figure 10). C2c2 cleaves RNA through conserved basic residues within its two HEPN domains, in contrast to the catalytic mechanisms of other known RNases found in CRISPR-Cas systems (Benda et al., 2014; Tamulaitis et al., 2014). Alanine substitution of any of the four predicted HEPN domain catalytic residues converted C2c2 into an inactive programmable RNA-binding protein (dC2c2, analogous to dCas9). Many different spacer sequences work well in our assays although further screening will likely define properties and rules governing optimal function.

These results suggest a broad range of biotechnology applications and research questions (Abil and Zhao, 2015; Filipovska and Rackham, 2011; Mackay et al., 2011). For example, the ability of dC2c2 to bind to specified sequences could be used to (i) bring effector modules to specific transcripts to modulate their function or translation, which could be used for large-scale screening, construction of synthetic regulatory circuits and other purposes; (ii) fluorescently tag specific RNAs to visualize their trafficking and/or localization; (iii) alter RNA localization through domains with affinity for specific subcellular compartments; and (iv) capture specific transcripts (through direct pull down of dC2c2) to enrich for proximal molecular partners, including RNAs and proteins.

Active C2c2 also has many potential applications such as targeting a specific transcript for destruction, as performed here with RFP. In addition, C2c2, once primed by the cognate target, can cleave other (non-complementary) RNA molecules *in vitro* and inhibit cell growth *in vivo.* Biologically, this promiscuous RNase activity might reflect a PCD/dormancy-based protection mechanism of the type VI CRISPR-Cas systems (Figure 7). Technologically, it might be used to trigger PCD or dormancy in specific cells such as cancer cells expressing a particular transcript, neurons of a given class, or cells infected by a specific pathogen.

Further experimental study is required to elucidate the mechanisms by which the C2c2 system acquires spacers and the classes of pathogens against which it protects bacteria. The presence of the conserved CRISPR adaptation module consisting of typical Cas1 and Cas2 proteins in the LshC2c2 locus suggests that it is capable of spacer acquisition. Although C2c2 systems lack reverse transcriptases, which mediate acquisition of RNA spacers in some type III systems (Silas et al., 2016), it is possible that additional host or viral factors could support RNA spacer acquisition. Additionally or alternatively, type VI systems could acquire DNA spacers similar to other CRISPR-Cas variants but then target transcripts of the respective DNA genomes, eliciting PCD and abortive infection (Figure 7).

The CRISPR-C2c2 system represent a distinct evolutionary path among Class 2 CRISPR-Cas systems. It is likely that other, broadly analogous Class 2 RNA-targeting immune systems exist, and further characterization of the diverse members of Class 2 systems will provide a deeper understanding of bacterial immunity and provide a rich starting point for the development of programmable molecular tools for *in vivo* RNA manipulation.

## Materials and Methods

Expanded materials and methods, including computational analysis, can be found in supplementary materials and methods.

## Bacterial phage interference

The C2c2 CRISPR locus was amplified from DNA from *Leptotrichia shahii DSM19757* (ATCC, Manassas, VA) and cloned for heterologous expression in *E. coli.* For screens, a library of all possible spacers targeting the MS2 genome were cloned into the spacer array; for individual spacers, single specific spacers were cloned into the array. Interference screens were performed in liquid culture and plated; surviving colonies were harvested for DNA and spacer representation was determined by next-generation sequencing. Individual spacers were tested by spotting on top agar.

## β-lactamase and transcribed/non-transcribed PFS preference screens

Sequences with randomized nucleotides adjacent to protospacer 1 were cloned into pUC19 in corresponding regions. Libraries were screened by co-transformation with LshC2c2 locus plasmid or pACYC184 plasmid control, harvesting of the surviving colonies, and next-generation sequencing of the resulting regions.

## RFP targeting assay

Cells containing an RFP expressing plasmid were transformed with an LshC2c2 locus plasmid with corresponding spacers, grown overnight, and analyzed for RFP fluorescence by flow cytometry. The growth effects of LshC2c2 activity were quantified by titrating inducible RFP levels with dilutions of anhydrotetracycline inducer and then measuring OD_600_.

## *in vitro* bnuclease and electrophoretic mobility shift assays

LshC2c2 protein and HEPN mutants were purified for use in *in vitro* reactions; RNA were synthesized via *in vitro* transcription. For nuclease assays, protein was co-incubated with crRNA and either 3’ or 5’-labeled targets and analyzed via denaturing gel electrophoresis and imaging or by next-generation sequencing. For electrophoretic mobility shift assays, protein and nucleic acid were co-incubated and then resolved by gel electrophoresis and imaging.

## Acknowledgements

We would like to thank P. Boutz, J. Doench, P. Sharp, and B. Zetsche for helpful discussions and insights; R. Belliveau for overall research support; J. Francis and D. O’Connell for generous MiSeq instrument access; D. Daniels and C. Garvie for providing bacterial incubation space for protein purification; R. Macrae for critical reading of the manuscript; and the entire Zhang laboratory for support and advice. We would like to thank N. Ranu for generously providing pRFP and D. Daniels for providing 6-His-MBP-TEV. O.A.A. is supported by a Paul and Daisy Soros Fellowship, a Friends of the McGovern Institute Fellowship, and the Poitras Center for Affective Disorders. J.S.G. is supported by a D.O.E. Computational Science Graduate Fellowship. S.S. is supported by the graduate program of Skoltech Data-Intensive Biomedicine and Biotechnology Center for Research, Education, and Innovation. I.S. is supported by the Simons Center for the Social Brain. D.B.T.C. is supported by award number T32GM007753 from the National Institute of General Medical Sciences. K.S.M., E.V.K. and, in part, S.S. are supported by the intramural program of the US Department of Health and Human services (to the National Library of Medicine). K.S. is supported by an NIH grant GM10407, Russian Science Foundation grant 14-14-00988, and Skoltech. F.Z. is a New York Stem Cell Foundation-Robertson Investigator. F.Z. is supported by the NIH through NIMH (5DP1-MH100706 and 1R01-MH110049), NSF, the New York Stem Cell, Simons, Paul G. Allen Family, and Vallee Foundations; and James and Patricia Poitras, Robert Metcalfe, and David Cheng. The authors plan to make the reagents widely available to the academic community through Addgene and to provide software tools via the Zhang lab website (www.genome-engineering.org)and GitHub (github.com/fengzhanglab).

## References

Abil, Z., and Zhao, H. (2015). Engineering reprogrammable RNA-binding proteins for study and manipulation of the transcriptome. Mol Biosyst 11, 2658–2665.

Anantharaman, V., Makarova, K.S., Burroughs, A.M., Koonin, E.V., and Aravind, L. (2013). Comprehensive analysis of the HEPN superfamily: identification of novel roles in intra-genomic conflicts, defense, pathogenesis and RNA processing. Biol Direct 8, 15.

Barrangou, R., Fremaux, C., Deveau, H., Richards, M., Boyaval, P., Moineau, S., Romero, D.A., and Horvath, P. (2007). CRISPR provides acquired resistance against viruses in prokaryotes. Science 315, 1709–1712.

Benda, C., Ebert, J., Scheltema, R.A., Schiller, H.B., Baumgartner, M., Bonneau, F., Mann, M., and Conti, E. (2014). Structural model of a CRISPR RNA-silencing complex reveals the RNA-target cleavage activity in Cmr4. Mol Cell 56, 43–54.

Brouns, S.J., Jore, M.M., Lundgren, M., Westra, E.R., Slijkhuis, R.J., Snijders, A.P., Dickman, M.J., Makarova, K.S., Koonin, E.V., and van der Oost, J. (2008). Small CRISPR RNAs guide antiviral defense in prokaryotes. Science 321, 960–964.

Crooks, G.E., Hon, G., Chandonia, J.M., and Brenner, S.E. (2004). WebLogo: a sequence logo generator. Genome research 14, 1188–1190.

Dahlman, J.E., Abudayyeh, O.O., Joung, J., Gootenberg, J.S., Zhang, F., and Konermann, S. (2015). Orthogonal gene knockout and activation with a catalytically active Cas9 nuclease. Nat Biotechnol 33, 1159–1161.

Datsenko, K.A., Pougach, K., Tikhonov, A., Wanner, B.L., Severinov, K., and Semenova, E. (2012). Molecular memory of prior infections activates the CRISPR/Cas adaptive bacterial immunity system. Nat Commun 3, 945.

Deltcheva, E., Chylinski, K., Sharma, C.M., Gonzales, K., Chao, Y., Pirzada, Z.A., Eckert, M.R., Vogel, J., and Charpentier, E. (2011). CRISPR RNA maturation by trans-encoded small RNA and host factor RNase III. Nature 471, 602–607.

Deng, L., Garrett, R.A., Shah, S.A., Peng, X., and She, Q. (2013). A novel interference mechanism by a type IIIB CRISPR-Cmr module in Sulfolobus. Mol Microbiol 87, 1088–1099.

Diez-Villasenor, C., Guzman, N.M., Almendros, C., Garcia-Martinez, J., and Mojica, F.J. (2013). CRISPR-spacer integration reporter plasmids reveal distinct genuine acquisition specificities among CRISPR-Cas I-E variants of Escherichia coli. RNA Biol 10, 792–802.

Filipovska, A., and Rackham, O. (2011). Designer RNA-binding proteins: New tools for manipulating the transcriptome. RNA Biol 8, 978–983.

Garneau, J.E., Dupuis, M.E., Villion, M., Romero, D.A., Barrangou, R., Boyaval, P., Fremaux, C., Horvath, P., Magadan, A.H., and Moineau, S. (2010). The CRISPR/Cas bacterial immune system cleaves bacteriophage and plasmid DNA. Nature 468, 67–71.

Gasiunas, G., Barrangou, R., Horvath, P., and Siksnys, V. (2012). Cas9-crRNA ribonucleoprotein complex mediates specific DNA cleavage for adaptive immunity in bacteria. Proc Natl Acad Sci U S A 109, E2579–2586.

Goldberg, G.W., Jiang, W., Bikard, D., and Marraffini, L.A. (2014). Conditional tolerance of temperate phages via transcription-dependent CRISPR-Cas targeting. Nature 514, 633–637.

Hale, C.R., Cocozaki, A., Li, H., Terns, R.M., and Terns, M.P. (2014). Target RNA capture and cleavage by the Cmr type III-B CRISPR-Cas effector complex. Genes Dev 28, 2432–2443.

Hale, C.R., Majumdar, S., Elmore, J., Pfister, N., Compton, M., Olson, S., Resch, A.M., Glover, C.V., 3rd, Graveley, B.R., Terns, R.M., et al. (2012). Essential features and rational design of CRISPR RNAs that function with the Cas RAMP module complex to cleave RNAs. Mol Cell 45, 292–302.

Hale, C.R., Zhao, P., Olson, S., Duff, M.O., Graveley, B.R., Wells, L., Terns, R.M., and Terns, M.P. (2009). RNA-guided RNA cleavage by a CRISPR RNA-Cas protein complex. Cell 139, 945–956.

Hayes, F., and Van Melderen, L. (2011). Toxins-antitoxins: diversity, evolution and function. Crit Rev Biochem Mol Biol 49, 386–408.

Heler, R., Samai, P., Modell, J.W., Weiner, C., Goldberg, G.W., Bikard, D., and Marraffini, L.A. (2015a). Cas9 specifies functional viral targets during CRISPR-Cas adaptation. Nature.

Heler, R., Samai, P., Modell, J.W., Weiner, C., Goldberg, G.W., Bikard, D., and Marraffini, L.A. (2015b). Cas9 specifies functional viral targets during CRISPR-Cas adaptation. Nature 519, 199–202.

Jackson, R.N., Golden, S.M., van Erp, P.B., Carter, J., Westra, E.R., Brouns, S.J., van der Oost, J., Terwilliger, T.C., Read, R.J., and Wiedenheft, B. (2014). Structural biology. Crystal structure of the CRISPR RNA-guided surveillance complex from Escherichia coli. Science 345, 1473–1479.

Jiang, W., Samai, P., and Marraffini, L.A. (2016). Degradation of Phage Transcripts by CRISPR-Associated RNases Enables Type III CRISPR-Cas Immunity. Cell 164, 710–721.

Jinek, M., Chylinski, K., Fonfara, I., Hauer, M., Doudna, J.A., and Charpentier, E. (2012). A programmable dual-RNA-guided DNA endonuclease in adaptive bacterial immunity. Science 337, 816–821.

Kiani, S., Chavez, A., Tuttle, M., Hall, R.N., Chari, R., Ter-Ovanesyan, D., Qian, J., Pruitt, B.W., Beal, J., Vora, S., et al. (2015). Cas9 gRNA engineering for genome editing, activation and repression. Nat Methods 12, 1051–1054.

Kim, Y.K., Kim, Y.G., and Oh, B.H. (2013). Crystal structure and nucleic acid-binding activity of the CRISPR-associated protein Csx1 of Pyrococcus furiosus. Proteins 81, 261–270.

Konermann, S., Brigham, M.D., Trevino, A.E., Joung, J., Abudayyeh, O.O., Barcena, C., Hsu, P.D., Habib, N., Gootenberg, J.S., Nishimasu, H., et al. (2015). Genome-scale transcriptional activation by an engineered CRISPR-Cas9 complex. Nature 517, 583–588.

Kozlov, G., Denisov, A.Y., Girard, M., Dicaire, M.J., Hamlin, J., McPherson, P.S., Brais, B., and Gehring, K. (2011). Structural Basis of Defects in the Sacsin HEPN Domain Responsible for Autosomal Recessive Spastic Ataxia of Charlevoix-Saguenay (ARSACS). J Biol Chem 286, 20407–20412.

Li, H., and Durbin, R. (2009). Fast and accurate short read alignment with Burrows-Wheeler transform. Bioinformatics 25, 1754–1760.

Liberzon, A., Birger, C., Thorvaldsdottir, H., Ghandi, M., Mesirov, J.P., and Tamayo, P. (2015). The Molecular Signatures Database (MSigDB) hallmark gene set collection. Cell Syst 1, 417–425.

Mackay, J.P., Font, J., and Segal, D.J. (2011). The prospects for designer single-stranded RNA-binding proteins. Nat Struct Mol Biol 18, 256–261.

Makarova, K.S., Anantharaman, V., Aravind, L., and Koonin, E.V. (2012). Live virus-free or die: coupling of antivirus immunity and programmed suicide or dormancy in prokaryotes. Biol Direct 7, 40.

Makarova, K.S., Haft, D.H., Barrangou, R., Brouns, S.J., Charpentier, E., Horvath, P., Moineau, S., Mojica, F.J., Wolf, Y.I., Yakunin, A.F., et al. (2011). Evolution and classification of the CRISPR-Cas systems. Nat Rev Microbiol 9, 467–477.

Makarova, K.S., Wolf, Y.I., Alkhnbashi, O.S., Costa, F., Shah, S.A., Saunders, S.J., Barrangou, R., Brouns, S.J., Charpentier, E., Haft, D.H., et al. (2015). An updated evolutionary classification of CRISPR-Cas systems. Nat Rev Microbiol 13, 722–736.

Makarova, K.S., Wolf, Y.I., and Koonin, E.V. (2009). Comprehensive comparative-genomic analysis of type 2 toxin-antitoxin systems and related mobile stress response systems in prokaryotes. Biol Direct 4, 19.

Marraffini, L.A., and Sontheimer, E.J. (2008). CRISPR interference limits horizontal gene transfer in staphylococci by targeting DNA. Science 322, 1843–1845.

Marraffini, L.A., and Sontheimer, E.J. (2010). Self versus non-self discrimination during CRISPR RNA-directed immunity. Nature 463, 568–571.

Mulepati, S., Heroux, A., and Bailey, S. (2014). Structural biology. Crystal structure of a CRISPR RNA-guided surveillance complex bound to a ssDNA target. Science 345, 1479–1484.

Niewoehner, O., and Jinek, M. (2016). Structural basis for the endoribonuclease activity of the type III-A CRISPR-associated protein Csm6. RNA 22, 318–329.

Nunez, J.K., Kranzusch, P.J., Noeske, J., Wright, A.V., Davies, C.W., and Doudna, J.A. (2014). Cas1-Cas2 complex formation mediates spacer acquisition during CRISPR-Cas adaptive immunity. Nature structural & molecular biology 21, 528–534.

Nunez, J.K., Lee, A.S., Engelman, A., and Doudna, J.A. (2015). Integrase-mediated spacer acquisition during CRISPR-Cas adaptive immunity. Nature.

Samai, P., Pyenson, N., Jiang, W., Goldberg, G.W., Hatoum-Aslan, A., and Marraffini, L.A. (2015). Co-transcriptional DNA and RNA Cleavage during Type III CRISPR-Cas Immunity. Cell 161, 1164–1174.

Sapranauskas, R., Gasiunas, G., Fremaux, C., Barrangou, R., Horvath, P., and Siksnys, V. (2011). The Streptococcus thermophilus CRISPR/Cas system provides immunity in Escherichia coli. Nucleic Acids Res 39, 9275–9282.

Sheppard, N.F., Glover, C.V., 3rd, Terns, R.M., and Terns, M.P. (2016). The CRISPR-associated Csx1 protein of Pyrococcus furiosus is an adenosine-specific endoribonuclease. RNA 22, 216–224.

Shmakov, S., Abudayyeh, O.O., Makarova, K.S., Wolf, Y.I., Gootenberg, J.S., Semenova, E., Minakhin, L., Joung, J., Konermann, S., Severinov, K., et al. (2015). Discovery and Functional Characterization of Diverse Class 2 CRISPR-Cas Systems. Mol Cell 60, 385–397.

Silas, S., Mohr, G., Sidote, D.J., Markham, L.M., Sanchez-Amat, A., Bhaya, D., Lambowitz, A.M., and Fire, A.Z. (2016). Direct CRISPR spacer acquisition from RNA by a natural reverse transcriptase-Cas1 fusion protein. Science 351, aad4234.

Sinkunas, T., Gasiunas, G., Waghmare, S.P., Dickman, M.J., Barrangou, R., Horvath, P., and Siksnys, V. (2013). In vitro reconstitution of Cascade-mediated CRISPR immunity in Streptococcus thermophilus. EMBO J 32, 385–394.

Staals, R.H., Agari, Y., Maki-Yonekura, S., Zhu, Y., Taylor, D.W., van Duijn, E., Barendregt, A., Vlot, M., Koehorst, J.J., Sakamoto, K., et al. (2013). Structure and activity of the RNA-targeting Type III-B CRISPR-Cas complex of Thermus thermophilus. Mol Cell 52, 135–145.

Staals, R.H., Zhu, Y., Taylor, D.W., Kornfeld, J.E., Sharma, K., Barendregt, A., Koehorst, J.J., Vlot, M., Neupane, N., Varossieau, K., et al. (2014). RNA targeting by the type III-A CRISPR-Cas Csm complex of Thermus thermophilus. Mol Cell 56 518–530.

Tamulaitis, G., Kazlauskiene, M., Manakova, E., Venclovas, C., Nwokeoji, A.O., Dickman, M.J., Horvath, P., and Siksnys, V. (2014). Programmable RNA shredding by the type III-A CRISPR-Cas system of Streptococcus thermophilus. Mol Cell 56, 506–517.

van der Oost, J., Jore, M.M., Westra, E.R., Lundgren, M., and Brouns, S.J. (2009). CRISPR-based adaptive and heritable immunity in prokaryotes. Trends Biochem Sci 34, 401–407.

Wright, A.V., Nunez, J.K., and Doudna, J.A. (2016). Biology and Applications of CRISPR Systems: Harnessing Nature's Toolbox for Genome Engineering. Cell 164, 29–44.

Yosef, I., Goren, M.G., and Qimron, U. (2012). Proteins and DNA elements essential for the CRISPR adaptation process in Escherichia coli. Nucleic Acids Res 40, 5569–5576.

Zetsche, B., Gootenberg, J.S., Abudayyeh, O.O., Slaymaker, I.M., Makarova, K.S., Essletzbichler, P., Volz, S.E., Joung, J., van der Oost, J., Regev, A., et al. (2015). Cpf1 is a single RNA-guided endonuclease of a class 2 CRISPR-Cas system. Cell 163, 759–771.

Zhang, J., Rouillon, C., Kerou, M., Reeks, J., Brugger, K., Graham, S., Reimann, J., Cannone, G., Liu, H., Albers, S.V., et al. (2012). Structure and mechanism of the CMR complex for CRISPR-mediated antiviral immunity. Mol Cell 45, 303–313.

